# Sex differences in insular gyri responses to the cold pressor challenge

**DOI:** 10.1101/2022.02.17.480949

**Authors:** Beatrix Krause-Sorio, Katherine E. Macey, Rajesh Kumar, Jennifer A. Ogren, Albert Thomas, Luke A. Henderson, Ronald M. Harper, Paul M. Macey

## Abstract

**Introduction:** Sex differences in autonomic control may contribute to physiological sex differences in cardiovascular disease, thermoregulation, and the experience of pain. We previously showed sex differences in functional mapping of autonomic responses across the insula, a central autonomic control region. Anterior insula response patterns to sympathetic activation (Valsalva) and parasympathetic withdrawal (handgrip) differed between men and women. We here assessed sex differences in autonomic insula gyrus responses to a cold pressor challenge, which involves temperature and pain regulation.

**Methods:** Functional MRI (fMRI) involved a 1-minute right-foot cold pressor challenge in 22 women (age; mean±std: 50±4 yrs), and 39 men (45±3 yrs). Regions of interest (ROI) comprised left and right anterior short gyrus (ASG), middle short gyrus (MSG), posterior short gyrus (PSG), anterior long gyrus (ALG), and posterior long gyrus (PLG). Two-second time intervals of fMRI signal responses per ROI and concurrently recorded heart rate and blood oxygen saturation (SaO_2_) were tested for within- and between-group effects over time (repeated measures ANOVA *P*≤0.05). We tested sex differences in 1) each ROI; 2) lateralization effects; and 3) posterior-to-anterior gradients in responses.

**Results:** No sex differences emerged in heart rate or SaO_2_. Women showed larger signal changes in several gyri than men after cold pressor onset and offset. Most consistently, women showed a greater right-over-left hemisphere dominance in cold pressor response compared to men in the MSG, PSG and ALG. Greater right-over-left hemisphere anterior-to-posterior dominance was more pronounced in men.

**Conclusions:** The findings confirm anterior and right-hemispheric dominance of insular responses during sympathetic activation that includes pain and cold. Although distinct transient male-female differences in fMRI responses appeared during the cold pressor, unlike the Valsalva and handgrip, no sex differences in ASG lateralization emerged, suggesting that the previously found sex differences may not solely relate to regulating blood pressure responses.

## Introduction

Women show different cardiovascular disease characteristics from men that may result from sex-specific autonomic regulation. For example, autonomic functions are influenced by sex-related factors, such as internal organ sizes and proportions, hormone levels, and variations in the central regulation of autonomic outflow [1-7, for review, see 8]. The autonomic nervous system (ANS) maintains homeostasis of the body based on processing signals from the organs and the environment. Whilst most of the basic neural circuitry responsible for actions of the ANS, such as the control of blood pressure, heart rate, gastric motility, pupillary diameter, and saliva production, are located within the brainstem, these brainstem sites are modulated by higher order brain centers, including cortical sites [9]. One brain region that appears to be critical in the integration of both autonomic function and sensory information is the insular cortex, a large cortical area in the fold of the Sylvian fissure. Strokes in this region disrupt the perception of taste, pain and body temperature in addition to blood pressure regulation [10–13]. The insula’s role in autonomic regulation is highlighted by findings that insular strokes are often associated with significantly higher plasma norepinephrine and epinephrine levels, as well as elevated blood pressure, and sympathetic/parasympathetic imbalance compared to non-insular stroke [14].

Anatomically, the insula projects to multiple brain regions that regulate autonomic outflow, including the hypothalamus and brainstem [15–22]. The anterior insula projects to limbic, paralimbic and inferior frontal regions, while the posterior insula connects with parietal and posterior temporal areas [23]. The insula’s functional neuroanatomy appears to follow the pattern of its cortical folding. The insula typically comprises five anatomically distinct gyri: the anterior short gyrus (ASG), the middle short gyrus (MSG), and the posterior short gyrus together form the anterior insula, while the anterior long gyrus (ALG) and the posterior long gyrus (PLG) form the posterior insula, although individual variation in the number of distinguishable gyri can occur [17]. These gyri show different functional roles. For instance, we showed earlier that ASG responses are larger than other gyri to challenges that raise blood pressure and increase sympathetic activation, such as the Valsalva maneuver and static handgrip [24]. The dorsal anterior insula also appears to show topographical representation of pain responses for different body parts [25]. However, it is unclear whether this functional anatomy extends to the regulation of pain and cold.

A challenge that involves both temperature and pain regulation in addition to raising blood pressure is the cold pressor, which involves immersion of an extremity in near-freezing cold water [26–31]. Autonomic responses to the cold pressor challenge are reproducible, and show variations associated with coronary risk, and whether a person will subsequently develop hypertension and cardiovascular disease [32–34]. We previously demonstrated that heart rate changes during a cold pressor challenge were accompanied by increased right anterior insula activity in healthy adolescents [35]. We also found that the individual gyri showed distinct activity patterns in healthy adults [24], more specifically, the right anterior short gyrus (ASG), the most anterior portion of the insula showing the greatest signal increase, and the posterior gyrus, the smallest. No significant signal increases could be observed in the left hemisphere gyri during the cold pressor.

The objective of the current study was to map the functional responses of the five insular gyri in men and women during a cold pressor challenge. Based on the evidence on sex differences in pain, thermoregulation, and physiological and neural responses in specific gyri of the insula to the handgrip and Valsalva challenges, we hypothesized that sex differences in the response of individual insular gyri during the cold pressor challenge are present. Based on earlier findings, we hypothesized that the right anterior insular gyri would 1) show sex differences in fMRI responses to the cold pressor, 2) respond more strongly than posterior gyri, and 3) respond more strongly in the right compared to the left hemisphere. The significance of such findings would be to suggest that the sex-specific responses of the insula relate to blood pressure increases independent of the specific nature of sensory or motor stimuli that elicited such changes.

## Methods

### Participants

We studied 61 healthy adults (age ± std: 47.2 ± 8.8 years, range: 31–66 years; 39 men, 22 women). All participants had no history of cerebrovascular disease, myocardial infarction, heart failure, neurological disorders, or mental illness, and were not taking cardiovascular or psychotropic medications. Participants were recruited from the Los Angeles area, and did not weigh more than 125 kg (275 lbs) or have any metallic or electronic implants; the latter two issues are MRI scanner contraindications. All participants provided written informed consent in accordance with the Declaration of Helsinki, and the research protocol was approved by the Institutional Review Board of UCLA. No participants were taking exogenous sex hormones (for example, oral contraceptive pills, hormone replacement therapy, or testosterone therapy).

### Cold pressor protocol

The protocol consisted of a 5-minute fMRI scanning period segmented into a 2-minute baseline, a 1-minute cold pressor, and a 2-minute recovery. Participants remained supine in the scanner with the right foot and lower leg bare throughout the procedure. Prior to the start of the protocol, a researcher positioned the participant’s right foot on the scanner bed with the knee bent, pointing upward. A container filled with ankle-height cold water (4.0 °C) and towel were placed nearby. Prior to the start of the fMRI scan, the researcher placed both hands on the sides of the lower leg, where they remained until the end of the 5-minute protocol. This was to avoid fMRI signal changes associated with touching or removing touch. At the 2-minute mark, a signal indicated to the researcher to lift and fully immerse the right foot into the ice water container. While one researcher lifted the foot off the bed, a second researcher placed the container under the foot and the first researcher then lowered the foot in the container. The researcher held the immersed foot in place for one minute. Upon the challenge termination signal, the researcher then lifted the foot out of the water and the second researcher removed the basin from the scanner bed and placed the foot on a towel. The participant remained still until the end of the scan.

### Physiological signals

Physiological data were recorded with an analog-to-digital acquisition system (instruNet INET-100B, GWI Instruments, Inc., Somerville, MA). Heart rate was derived from the pulse waveforms of an MRI-compatible pulse oximeter (Nonin 8600FO, Nonin Medical Inc., Plymouth, MN). The pulse sensor was placed on the right index finger and HR was computed from the raw oximetry signal acquired at 1 kHz using custom peak-detection software followed by a careful visual review by an experienced researcher. All signals were synchronized with the MRI scans, and data corresponding to the fMRI recording period extracted. The HR value corresponding to each MRI image was derived through interpolation of the continuous beat-to-beat HR.

### MRI scanning

Functional MRI scans were acquired while participants lay supine using a 3.0-Tesla scanner (Siemens Magneton Tim-Trio, Erlangen, Germany), with a third party 8-channel head coil. A foam pad was placed on either side of the head to minimize movement. We collected whole-brain images with the blood-oxygen level dependent (BOLD) contrast (repetition time [TR] = 2000 ms; echo time [TE] = 30 ms; flip angle = 90°; matrix size = 64 × 64; field-of-view = 220 × 220 mm; slice thickness = 4.5 mm). The spatial resolution was based on achieving whole-brain coverage with the fastest possible acquisition time. Two high-resolution, T1-weighted anatomical images were also acquired with a magnetization prepared rapid acquisition gradient echo sequence (TR = 2200 ms; TE = 2.2 ms; inversion time = 900 ms; flip angle = 9°; matrix size = 256 × 256; field-of-view = 230 × 230 mm; slice thickness = 1.0 mm). Field map data consisting of phase and magnitude images were collected to allow for correction of distortions due to field inhomogeneities.

### MRI data preprocessing

All anatomical scans were inspected for visible pathology. For each fMRI series, the global signal was calculated, and the images realigned to account for head motion. Participants with large changes in global BOLD signal, or who moved more than 4 mm in any direction, were not included in the study. Each fMRI series was linearly detrended to account for signal drift [36], corrected for field inhomogeneities, spatially normalized, and across all participants. A mean image of all participants’ spatially normalized, anatomical scans was created. Software used included the statistical parametric mapping package, SPM12 (Wellcome Department of Cognitive Neurology, UK; www.fil.ion.ucl.ac.uk/spm), MRIcron [37], and MATLAB-based custom software.

### Region of interest (ROI)

The five major gyral regions in the insular cortex, the anterior short gyrus (ASG), mid short gyrus (MSG), posterior short gyrus (PSG), anterior long gyrus (ALG) and posterior long gyrus (PLG), were outlined on the mean anatomical image with MRIcroN software [37], using previously-published anatomical descriptions [17, 38]. Figure 1 illustrates the gryi on an average anatomical scan in sagittal and axial views. While individual tracing would be more accurate for identifying the gyral differentiation on anatomical scans, the lower spatial resolution of the fMRI images (voxel volume of 53 mm^3^ versus 0.8 mm^3^ for the anatomical scans) and the diffuse nature of the BOLD response meant that the advantage of individual tracing would be negligible. Due to individual variability in insular gyral folding [39], the present approach distinguishes gyral regions rather than gyri per se. The three main gyri of the anterior insula, the ASG, MSG, and PSG, make up the convex surface of the structure, and are visible on the sagittal and axial views of the mean anatomical image. The accessory and transverse gyri, two other gyri in the anterior insula, are difficult to visualize [38], and were not visible on the mean anatomical image. Thus, in our tracing of the ASG, we included the entire most-anterior portion of the insula, which included the accessory and transverse gyri. The posterior gyri (ALG and PLG) were easily visible on sagittal, as well as axial sections of the anatomical volume.

**Figure 1.**
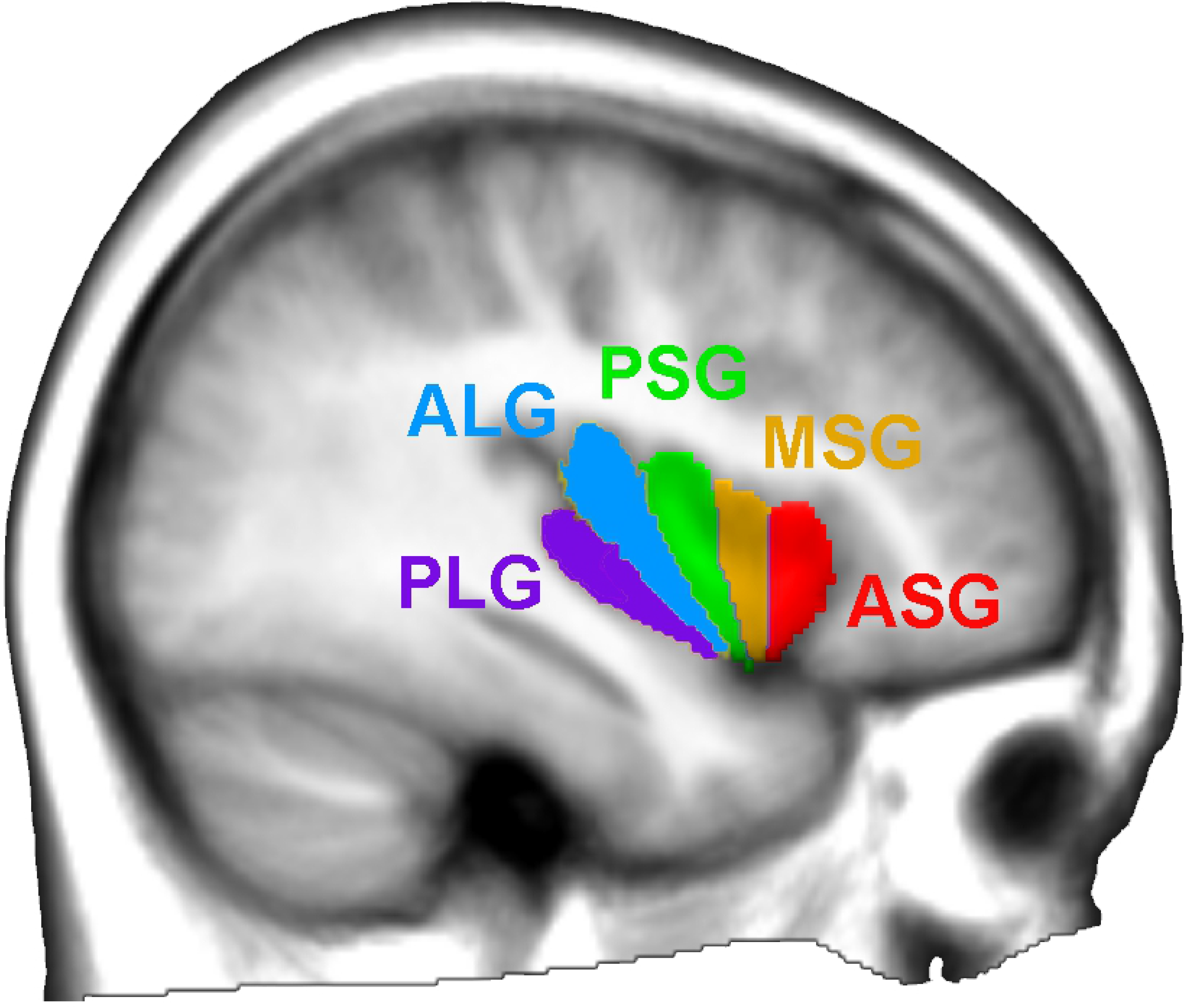
The five insular gyri color-coded and overlaid on the average of all participants’ anatomical scans normalized to template space. The anterior region of the insula is comprised of the short gyri, including the anterior short gyrus (ASG), mid short gyrus (MSG), and posterior short gyrus (PSG). The posterior region of the insula is comprised of the long gyri, including the anterior long gyrus (ALG) and posterior long gyrus (PLG).

### Statistical analysis

Demographic variables were assessed for group differences with chi-square and independent samples t-tests. Neural and HR responses were assessed by analysis of time trends. Repeated measures ANOVA (RMANOVA), implemented with the mixed linear model procedure “proc mixed” in SAS 9.4 software [40], was used to identify periods of significantly increased BOLD response relative to baseline, during the cold pressor challenge and subsequent recovery period; the RMANOVA approach is presented in detail elsewhere [41], including as used for heart rate assessments [42]. The restriction of only assessing differences rather than absolute responses results from the relative nature of the BOLD-based fMRI technique. We modeled the BOLD responses with group (i.e., gyrus or sex), time and group × time. For the “time” variable, scans in the 2 min baseline were categorized as “0”, and subsequent scans during the challenge and recovery were categorized sequentially (so, for example, the time “10” was the 10^th^ scan of the cold pressor challenge, occurring at 20 seconds after onset). Model significance was first assessed at the overall level; as per the Tukey-Fisher criterion for multiple comparisons, only models showing overall significance (*P* ≤ 0.05) were further assessed. Within-group responses (i.e., individual gyral responses from baseline during and after the challenge) were assessed for models with significant effects of time. Between-group responses (i.e., differences in responses between gyri) were assessed for models with significance effects of group × time. To identify anterior-posterior organization, we assessed responses with respect to the PLG, as the posterior insula typically responds less than anterior regions in response to autonomic stimuli [43]. Thus, we assessed in one model time trends ASG-PLG, AMG-PLG, PSG-PLG and ALG-PLG. To identify lateral organization, we assessed right-sided relative to left-sided responses for each gyrus [24]. We did not include a hemisphere factor in any model, since the aims were restricted to identifying gyral-specific differences. Three sets of models were created: 1) sex differences with a model of sex, time and sex × time for all gyri (10 models); 2) anterior-posterior differences with a model of gyri, time and gyri × time for each hemisphere and each sex, where gyri includes four categories: ASG-PLG, MSG-PLG, PSG-PLG and ALG-PLG (4 models); and 3) sex differences in laterality with a model of sex, time and sex × time for each bilateral gyrus, so ASG right – left, MSG right – left, PSG right – left, ALG right – left, and PLG right – left (5 models).

## Results

### Participants

There was no significant difference between women and men with respect to age (mean age ± std: women 50.0 ± 7.9 yrs, men 45.6 ± 9.0 yrs; *P* = 0.06, independent samples t-test), body mass index (BMI) (women 24.0 ± 5.1 kg/m^2^, men 25.1 ± 2.8 kg/m^2^; group difference *P* = 0.3, independent samples t-test), or handedness (6 women left-handed, 3 ambidextrous, 13 right-handed; 4 men left-handed, 2 ambidextrous, 33 right-handed,; *P* = 0.14, chi-squared).

### Physiology

HR showed significant time, but not sex or sex by time effects (time: *P* <.0001, sex: *P* = 0.07, sex × time: *P* = 0.7; 1 sec time bins for RMANOVA), with relative increases in both women and men during the cold period (model statistics: chi-squared = 1935, −2 res log likelihood = 127,565). The absolute HRs were higher in women than men (Figure 2). Oxygen saturation also only showed an effect of time (time: *P* <.0001, sex: *P* = 0.4, sex × time: *P* = 0.9).

**Figure 2.**
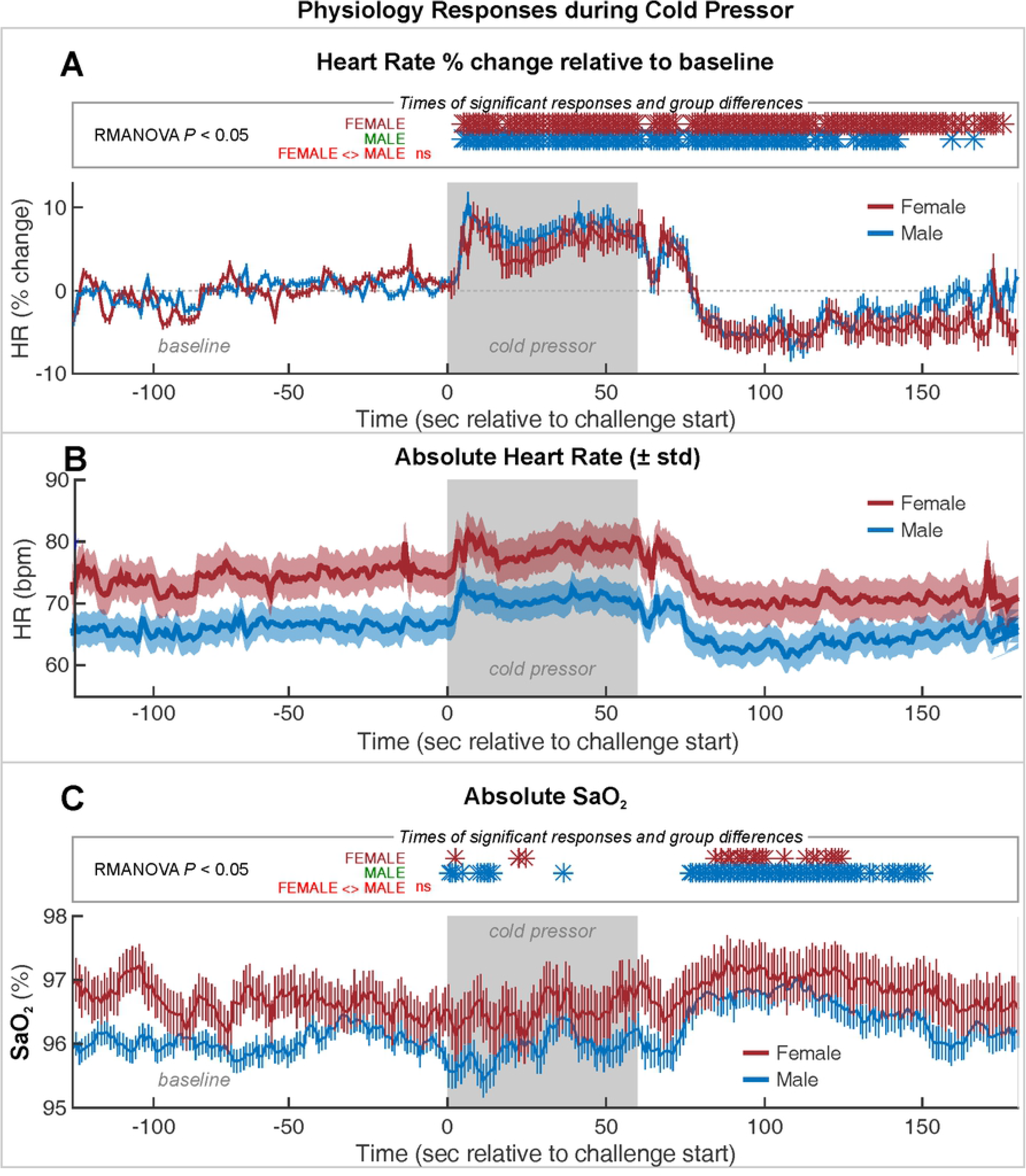
Average heart rate (HR) and SaO_2_ time course during baseline, cold pressor and recovery periods for men and women. **A)** HR percent change relative to 2 min baseline; **B)** absolute heart rate, showing a significantly higher baseline HR in females vs. males; and **C)** SaO_2_. Shading indicates SEM based on mean and error bars indicate EEM based on RMANOVA. Time points of significant within-group increases are indicated by blue and green asterisks in the top panel of each sub-figure; there were no significant between-group differences in responses (RMANOVA, *P* ≤ 0.05).

### BOLD response

The cold pressor elicited fMRI signal responses that differed from baseline during the one-minute cold period, with responses seen in all gyri across both sexes (within-group effects; Figure 3, Table 1). Note that the signal immediately after the cold pressor onset is lost on the y-axis; this drop is due to the passive movement of the foot into the water, which slightly shifted the body. The signal returns after 2-3 volumes. The lost signal is apparent only in Figure 3, as other figures involve relative signals.

**Figure 3.**
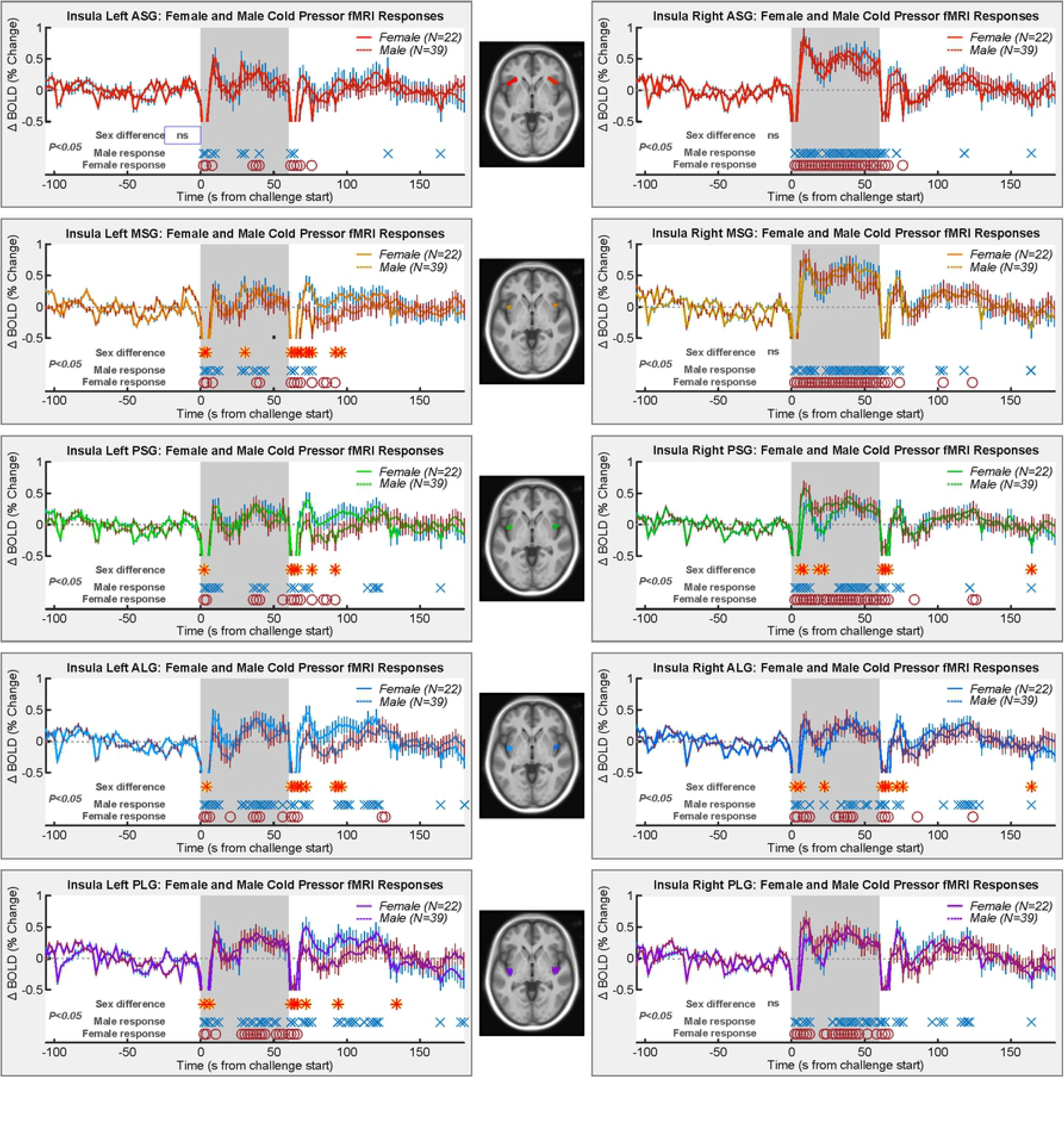
Mean insular gyri BOLD responses to the cold pressor in men and women. Time points of significant within-group responses and between group differences are indicated above the x-axis, and below the graphs (RMANOVA *P* ≤ 0.05; Table 1). Note that the initial drop of the signal after the cold pressor onset is cut off along the y-axis; this drop is due to the passive movement of the foot into the water bucket, therefore slightly shifting the body. This is apparent in the short duration of this shar signal drop, as the signal returns after only 2-3 volumes.

**Table 1.**
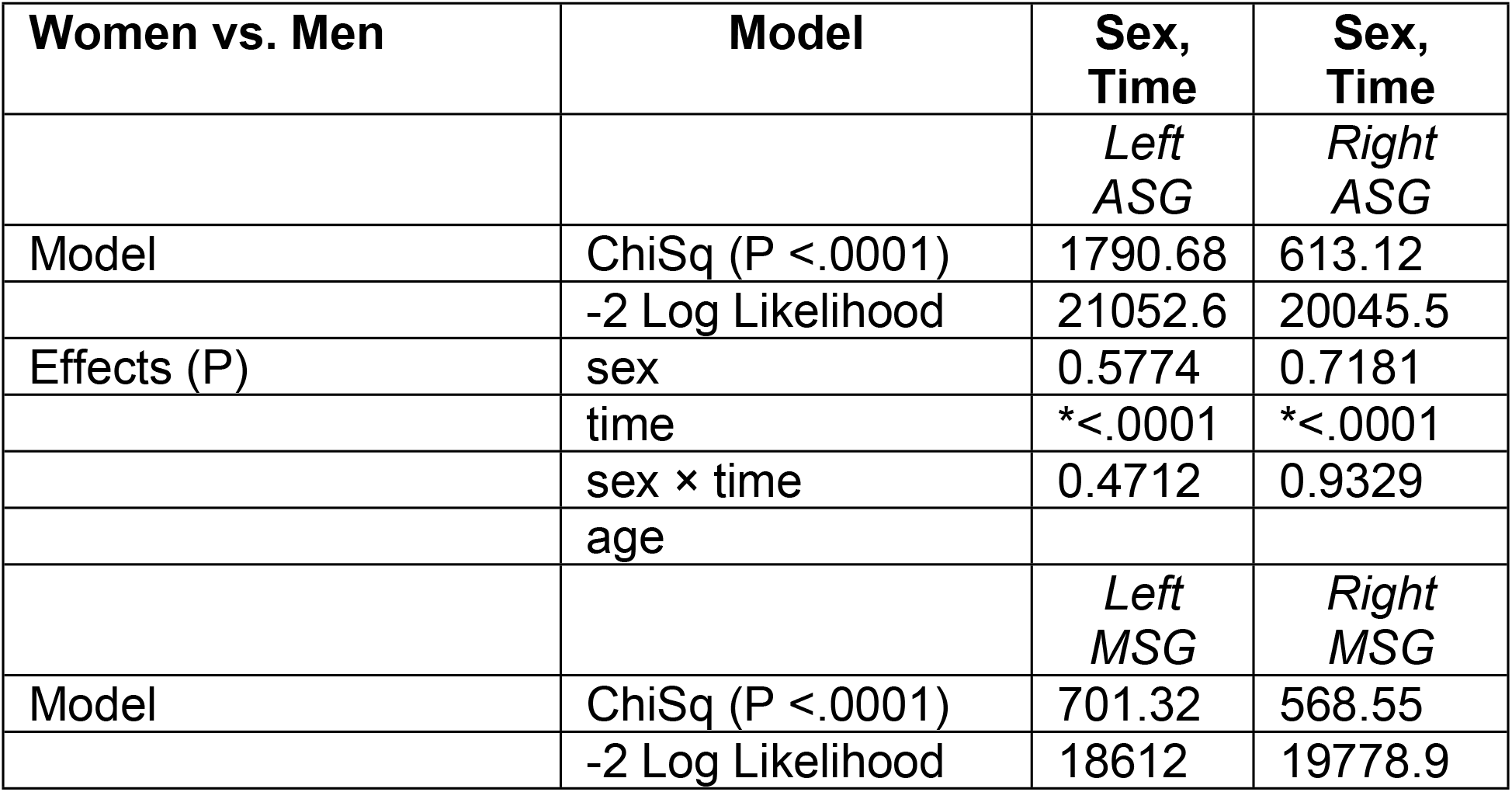
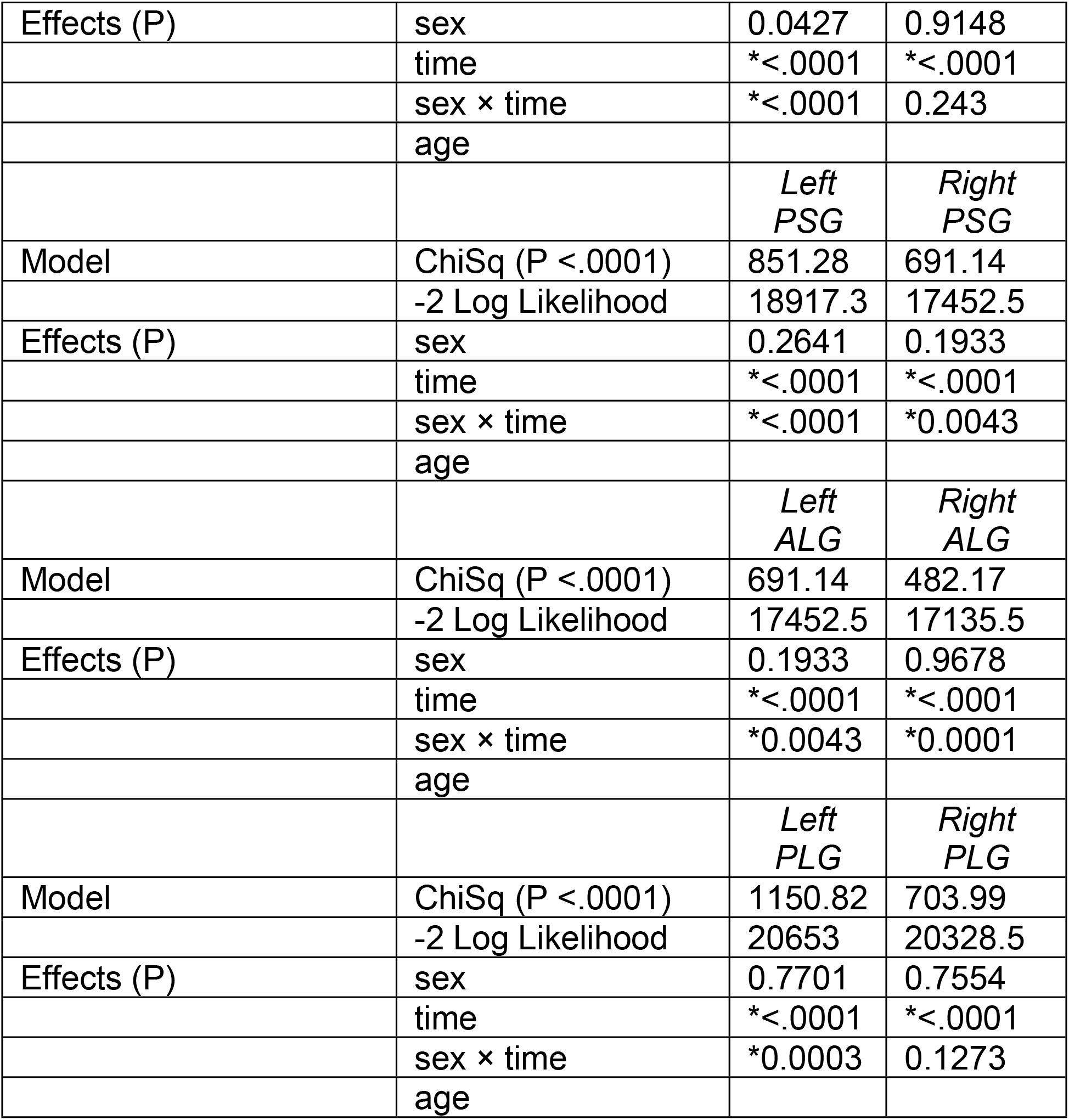
Model fit for women vs. men. The model Chi Square (ChiSq) was significant in both cases (P<0.0001). The model fit is indicated by −2 x log likelihood as calculated by SAS (higher indicates better fit). The *P*-values for each variable are shown; the sex × time interaction was consistently significant and age was not a significant covariate in any model.

| Women vs. Men | Model | Sex,<br>Time | Sex,<br>Time |
| --- | --- | --- | --- |
|  |  | *Left*<br>*ASG* | *Right*<br>*ASG* |
| Model | ChiSq ($P < .0001$) | 1790.68 | 613.12 |
|  | -2 Log Likelihood | 21052.6 | 20045.5 |
| Effects (P) | sex | 0.5774 | 0.7181 |
| | time | *$<.0001$ | *$<.0001$ |
| | sex × time | 0.4712 | 0.9329 |
|  | age |  |  |
|  |  | *Left*<br>*MSG* | *Right*<br>*MSG* |
| Model | ChiSq ($P < .0001$) | 701.32 | 568.55 |
|  | -2 Log Likelihood | 18612 | 19778.9 |
| Effects (P) | sex | 0.0427 | 0.9148 |
|  | time | *<.0001 | *<.0001 |
|  | sex × time | *<.0001 | 0.243 |
|  | age |  |  |
|  |  | *Left*<br>*PSG* | *Right*<br>*PSG* |
| Model | ChiSq (P <.0001) | 851.28 | 691.14 |
|  | -2 Log Likelihood | 18917.3 | 17452.5 |
| Effects (P) | sex | 0.2641 | 0.1933 |
|  | time | *<.0001 | *<.0001 |
|  | sex × time | *<.0001 | *0.0043 |
|  | age |  |  |
|  |  | *Left*<br>*ALG* | *Right*<br>*ALG* |
| Model | ChiSq (P <.0001) | 691.14 | 482.17 |
|  | -2 Log Likelihood | 17452.5 | 17135.5 |
| Effects (P) | sex | 0.1933 | 0.9678 |
|  | time | *<.0001 | *<.0001 |
|  | sex × time | *0.0043 | *0.0001 |
|  | age |  |  |
|  |  | *Left*<br>*PLG* | *Right*<br>*PLG* |
| Model | ChiSq (P <.0001) | 1150.82 | 703.99 |
|  | -2 Log Likelihood | 20653 | 20328.5 |
| Effects (P) | sex | 0.7701 | 0.7554 |
|  | time | *<.0001 | *<.0001 |
|  | sex × time | *0.0003 | 0.1273 |
|  | age |  |  |

For men and women, significant within-group responses were present throughout the cold period and during the recovery (men: blue X’s and women: red O’s in Figure 3). A consistent overall pattern of fMRI signal response to the cold pressor was evident in all ROIs of both the right and left insula. At the onset of the cold pressor period, there was an initial transient decrease in signal followed by a rapid recovery and increase above baseline. This increase then either dipped and then increased or remained constant above baseline for the remainder of the stimulus period. At the completion of the cold pressor period, signal intensity again transiently dipped below baseline before increasing to baseline, where it remained. Whilst there were no significant differences in fMRI signal changes between men and women in the right anterior/mid (ASG, MSG) and posterior insula (PLG) regions, there were significant differences in the mid/posterior insular gyri. In the right PSG and ALG regions, the transient signal decreases at the onset and immediately following the end of the cold pressor periods were greater in magnitude in women compared with men.

Overall, the patterns of signal changes were similar between the left and right insula, while overall signal changes in the left insula were smaller in magnitude, particularly within more anterior regions. However, like the right insula, the transient signal declined at the start and immediately following the end of the cold pressor periods were greater in magnitude in women compared with men in all regions apart from the very anterior (ASG) insula.

Although there were no significant differences in overall signal changes through the cold pressor task in men compared with women, examination of the relative contributions of anterior versus posterior regions revealed some sex differences (Figure 4, Table 2). Compared to the PLG, in men, the right anterior gyri showed greater responses to this sympathetic challenge with the ASG and MSG showing differences between all the posterior gyri. The left gyri showed no anterior-posterior response differences in men. However, in direct contrast to men, in women, there were no anterior-posterior response differences on either the left or the right insula gyri.

**Figure 4.**
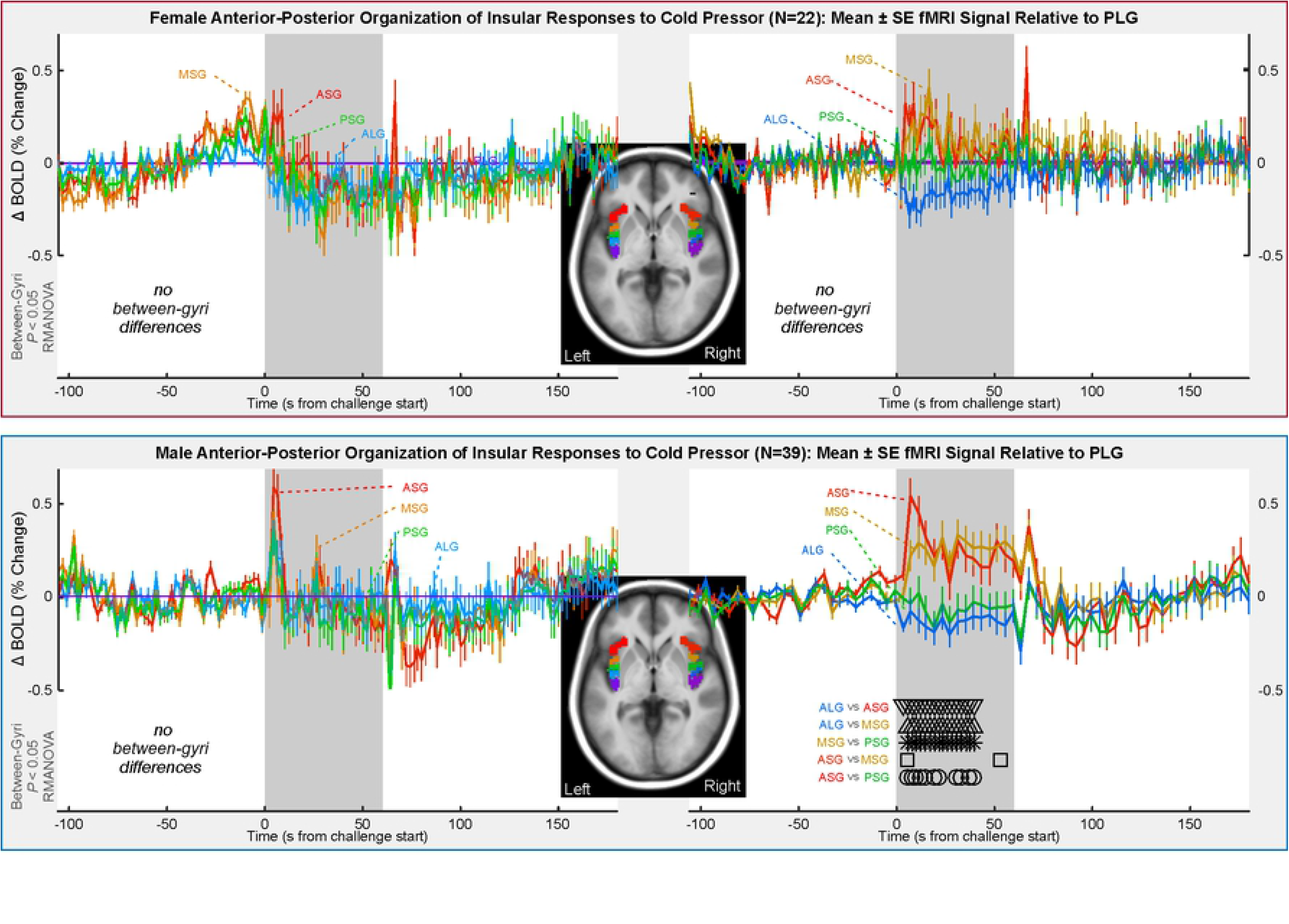
Anterior-to-posterior organization of insular gyri BOLD responses relative to the most posterior gyrus (PLG). Signals for women are shown in the top and men in the bottom panel. Time points of between-gyrus differences are indicated by symbols above the x-axis and below the graphs (RMANOVA *P* ≤ 0.05; Table 2).

**Table 2.** Anterior-posterior model fit. Overall model Chi Square (ChiSq) was consistently significant (P<0.0001). The model fit is indicated by −2 x log likelihood as calculated by SAS (higher indicates better fit). The *P*-values for each variable are shown; the gyrus × time interaction was consistently significant.

| Anterior vs. Posterior Gyri |  |  |  |
| --- | --- | --- | --- |
| Model |  | Gyrus, Time | Gyrus, Time |
|  | ChiSq (P <.0001) | 1723.92 | 1421.33 |
|  | -2 Log Likelihood | 16168 | 14372.7 |
| Effects (P) | gyrus | 0.7488 | 0.0011 |
|  | time | <.0001 | <.0001 |
|  | gyrus × time | 1 | 0.2033 |
|  | age |  |  |
| Model |  | Gyrus, Time | Gyrus, Time |
|  | ChiSq (P <.0001) | 1735.71 | 1150.3 |
|  | -2 Log Likelihood | 45614.9 | 16975.2 |
| Effects (P) | gyrus | 0.9214 | 0.0002 |
|  | time | <.0001 | <.0001 |
|  | gyrus × time | 1 | <.0001 |
|  | age |  |  |

Assessment of lateralization of activity during the challenge revealed significantly greater signal intensity changes on the right side in the anterior/mid insula (ASG, MSG) for both men and women. In women, the PSG and PLG also showed greater right sided signal increases during the first half of the cold pressor challenge period (Table 3; Figure 5). In addition, in the MSG and PSG, women showed a greater lateralization than men for the first 30 seconds of the challenge. In the ALG, there was no significant effect.

**Figure 5.**
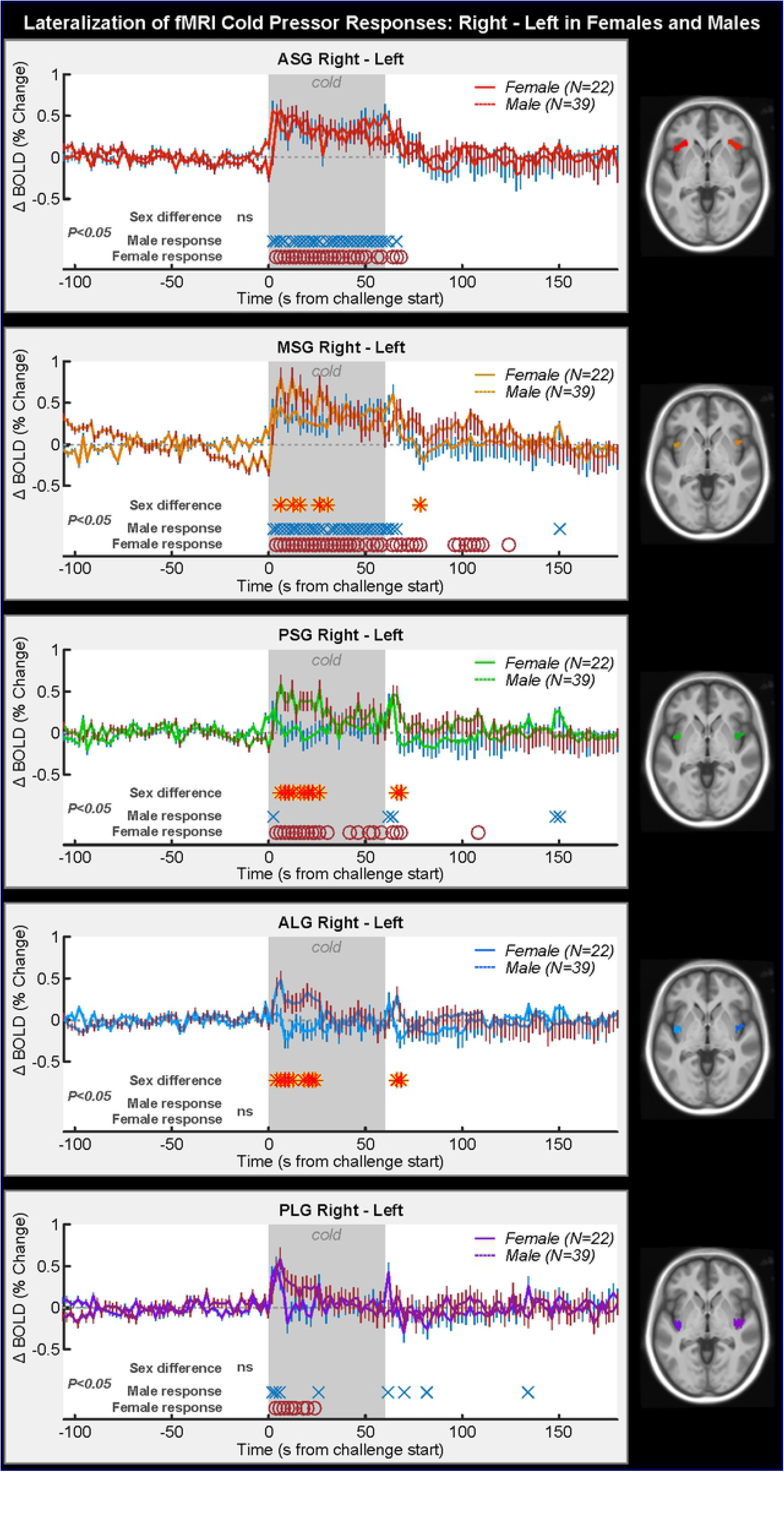
Lateralization of insular gyri BOLD responses in females and males. The signal of the left hemisphere gyri was subtracted from the right hemisphere gyri for each insular region, such that a higher signal indicates a greater right-lateralized response. Time points of between-hemisphere differences in women and men are indicated, as well as time points of group differences (RMANOVA *P* ≤ 0.05; Table 3).

**Table 3.** Left vs. right model fit. Overall model Chi Square (ChiSq) was always significant (P<0.0001). The model fit is indicated by −2 x log likelihood as calculated by SAS (higher indicates better fit). The *P* values for each variable are shown; sex × time interaction and time were significant (*P* < 0.05) in all cases apart from the ASG in men and MSG in women.

| Right vs. Left | Model | Sex, Time |
| --- | --- | --- |
| **ASG** |  |  |
| Model | ChiSq (P <.0001) | 1502.54 |
|  | -2 Log Likelihood | 17306.5 |
| Effects (P) | sex | 0.7839 |
|  | time | *<.0001 |
|  | sex × time | 0.6883 |
|  | age |  |
| **MSG** |  |  |
| Model | ChiSq (P <.0001) | 804.57 |
|  | -2 Log Likelihood | 17429.4 |
| Effects (P) | sex | 0.1056 |
|  | time | *<.0001 |
|  | sex × time | *0.0025 |
|  | age |  |
| **PSG** |  |  |
| Model | ChiSq (P <.0001) | 1341.81 |
|  | -2 Log Likelihood | 17408.1 |
| Effects (P) | sex | 0.0606 |
|  | time | *<.0001 |
|  | sex × time | *0.0008 |
|  | age |  |
| **ALG** |  |  |
| Model | ChiSq (P <.0001) | 954.71 |
|  | -2 Log Likelihood | 15710.3 |
| Effects (P) | sex | 0.1341 |
|  | time | 0.5048 |
|  | sex × time | *0.0134 |
|  | age |  |
| **PLG** |  |  |
| Model | ChiSq (P <.0001) | 1023.56 |
|  | -2 Log Likelihood | 17233.5 |
| Effects (P) | sex | 0.6069 |
|  | time | *<.0001 |
|  | sex × time | 0.6747 |
|  | age |  |

## Discussion

During a cold pressor challenge, HR increased from baseline by a similar percentage in both men and women. The challenge elicited increases in insular fMRI signal intensity over the entire challenge period that followed a similar pattern in men and women, and were primarily right-lateralized. Despite similarities in timing, the magnitude of transient dips in signal intensity at the onset and immediately after the end of the challenge were significantly greater in women compared to men. Furthermore, analysis of anterior-posterior insular responses revealed significantly greater signal intensity changes in the right anterior insula in men but no differences in women. This pattern contrasted with our previous findings during a Valsalva maneuver, in which we reported stronger anterior insula responses in women compared to men [44]. Finally, lateralization effects revealed greater signal intensity changes in right-lateralized anterior/mid insula for both men and women. This ipsilateral right-insula activation occurred despite the what would be expected for sensory stimulation of the right foot. We contrasted the insular findings from the present cold pressor and previously published Valsalva and hand grip challenges in Table 4.

**Table 4.** Overview of the task demands and results from the current study using the cold pressor challenge contrasted with our previous work using the Valsalva maneuver and the static hand grip challenge [44, 45].

| Effects | Cold pressor | Valsalva | Handgrip |
| --- | --- | --- | --- |
| | Foot is placed in 4°C cold water for one minute | The participant blows into a tube at $\geq 30$mmHg for 18 seconds | The participant squeezes a hard rubber ball for 16 seconds at a static 80% of their maximum grip strength |
| ANS function | Sympathetic activation and parasympathetic withdrawal | Sympathetic activation | Parasympathetic withdrawal, most likely vagal withdrawal |
| Organ effect | Vasoconstriction of the blood vessels | Thoracic pressure from expansion of the lungs | Cardiovascular pressure response due to muscle activity (breathing by itself is not altered) |
| Cardiovascular | Vasoconstriction | Thoracic pressure exerts pressure on heart, therefore higher pump capacity | Vasoconstriction |
| Observed heart rate changes | Increased heart rate and SaO<sub>2</sub> during stimulation | Increased heart rate | Increased heart rate |
| Insula gyri | Right ASG and MSG most prominent responses | Left hemisphere between-gyrus differences:<br>Men: ASG>MSG>PSG>PLG<br>Women: ASG≥MSG>PSG≥PLG<br>Right hemisphere between-gyrus differences:<br>Men:<br>ASG/MSG>PSG>ALG/PLG<br>Women:<br>MSG/PSG/PLG>ASG/ALG | Right ASG<br>The right anterior insula showed stronger responses than the posterior across men and women and there were no differences between the posterior three gyri. |
| Lateralization | Stronger right-lateralized responses in ASG and MSG, less so in the PLG<br>No lateralization effect in the ALG<br>Women showed an increased lateralization in the PSG but men did not | Stronger right-lateralized responses in all gyri, except in the ASG of women. | Posterior gyri showed stronger activation in the left compared to the right hemisphere |
| Sex differences | Heart rate and SaO<sub>2</sub> showed no sex differences but women had a higher variability in the SaO<sub>2</sub> signal at baseline, which may be responsible for the lack of effects.<br>Men showed significant decreases in SaO<sub>2</sub> at the beginning of the cold pressor and the largest | Larger heart rate increases in women than men, but smaller relative changes.<br>Women showed slightly higher SaO<sub>2</sub> at baseline and recovery compared to men, and a larger decline in SaO<sub>2</sub> during the challenge.<br>Women showed consistently higher responses than men in all gyri of both hemispheres (men showed sharper signal declines).<br>The right ASG was the only gyrus that showed no sex differences during a small period 13-15 seconds into the Valsalva.<br>Lateralization effects were observable in all gyri except in | Women showed smaller heart rate increases than men despite higher resting heart rate<br>SaO<sub>2</sub> decreased in men but not women and remained low throughout the recovery period and subsequent handgrip trials.<br>Women showed higher insula responses than men in all gyri |
|  | part of the recovery period. There were no sex differences in left or right ASG or the right MSG or PLG. Sex differences in other gyri were limited to the first peak of the BOLD response after cold pressor onset and the sharp drop in BOLD response after the offset of stimulation. Women showed higher right-over-left dominance in the MSG, PLG and ALG during the first third of the cold pressor stimulation compared to men. | the ASG in men and the MSG in women. | except the left ASG. Women showed greater right-over-left activation, while men showed the opposite pattern. Only men showed stronger left-lateralized increases in the middle gyrus |

The fMRI signal changes in the insula are likely neural activity reflecting a combination of salience of the stimulus of the foot being placed in cold water and of the regulation of the autonomic responses to the pressor challenge [46]. We focused on autonomic responses and sex differences in cardiovascular responses elicited by the cold pressor, as demonstrated in earlier studies. For example, older women with elevated blood pressure showed higher blood pressure increases in response to the cold pressor compared to younger women and men, while postmenopausal women showed stronger sympathetic responses to the cold pressor compared to age-matched men [47, 48]. In another study, compared to men, women show higher heart rate, lower systolic and diastolic blood pressure, and reduced autonomic responses (parasympathetic nervous system indicator defined as heart rate variability high frequency/total power) in response to the cold pressor [31]. A meta-analysis of twin studies provided strong evidence that the heart rate and blood pressure responses to the cold pressor occur differently in men and women [49]. In a study focused on obstructive sleep apnea, our group showed in healthy people the presence of sex differences in heart rate changes to the cold pressor [42]. Thus, we expected that underlying these cardiovascular sex differences we would observe neural sex differences in insula fMRI responses.

Women displayed greater transient fMRI signal intensity changes than men at onset and after challenge cessation in both hemispheres, predominantly in the mid/posterior insula regions. We previously showed that heart rate changes during a cold pressor challenge were accompanied by increased right anterior insula activity in healthy adolescents [35]. We also found that the individual gyri showed distinct activity patterns in healthy adults [24], more specifically, the right anterior short gyrus (ASG), the most anterior portion of the insula showing the greatest signal increase and the posterior gyrus, the smallest.

No significant signal increases could be observed in the gyri of the left hemisphere during the cold pressor. However, women showed significantly higher right-over-left dominance in the MSG, PLG and ALG during the first third of the cold pressor stimulation compared to men. Only in men, right but not left ASG and MSG showed significantly increased BOLD responses compared to the posterior gyri throughout the cold pressor. It is important to emphasize again that the anterior-posterior analysis was based on the difference between each gyrus and the PLG. The individual BOLD response pattern still demonstrated a gradually increasing cold pressor response from posterior to anterior gyri in both men and women. However, this specific result in men emphasizes the dominant role of ASG and MSG in this physiological challenge, which co-occurs with a slightly, even though not significantly higher change in heart rate than in women.

Other challenges elicit sex differences in insular responses; earlier, we found greater insula deactivation to the Valsalva maneuver in men compared to women [44] and hand grip responses in the anterior insular gyri were greater in the right hemisphere in women and greater in the left hemisphere in men [45]. Other groups have found opposite responses whereby a hand grip task led to higher right-sided responses in men compared to women [50]. Furthermore, lower body negative pressure led to greater right, but not left insular activity in men compared with women [51, 52].

The uniqueness of the sex differences to the cold pressor compared with handgrip and Valsalva likely relates to the cold and pain aspect of the challenge, as opposed to the pressor component present in all three challenges. One of the most consistent findings with respect to insula function is its involvement in the detection of novel stimuli across different sensory modalities, and in particular, temperature and noxious stimuli. The insula, together with the anterior cingulate cortex and other subcortical structures are referred to as the salience network, and are critical in identifying the most homeostatically relevant competing internal and external stimuli [53]. The insula is typically cited as a region that processes and integrates internal body sensations [54], i.e. interoception, including body temperature [55]. Interestingly, activity in the posterior insula correlates with changes in objective thermal intensity, whereas activity in the right anterior insula correlates with perceived thermal intensity [46]. Furthermore, the posterior insula is involved in innocuous-temperature perception and the anterior insula the perception of pain [56, 57]. These reports are seemingly consistent with our findings in that the sharp change in signal intensity in the left and right posterior/mid insula when the foot is being placed into and out of the cold water, likely reflects both salience as well as the physical detection of a temperature change in the foot. In contrast, the sustained increase in signal intensity almost restricted to the right anterior insula reflects the prolonged unpleasant and painful aspects of the cold pressor challenge.

Given potential general sex differences in fMRI responses, sex differences in the functional state of the insula are likely present, perhaps due to structural or hormonal differences. For example, women show greater gray matter volume in the right insular cortex than men [58], while on average, men show a greater number of distinct anatomical insular gyri [4.7 vs. 4.1; 39]. Specifically, the MSG is more clearly distinguishable in men, particularly in the left hemisphere, suggesting sex-and hemisphere-related regional specificity in insular anatomy [59]. Sex differences are also found in insular structural connectivity and metabolism at a younger age [60, 61]. Sex hormones affect various brain functions [62–64], and can have neuroprotective effects [65]. Sex differences are found in regional steroid receptor concentrations and distributions of major autonomic network components (e.g., the insula, amygdala, and anterior cingulate gyrus) [58, 62, 63]. Furthermore, global cerebral blood flow (CBF) is higher in women than men, and activity-related CBF changes vary by sex as well [66, 67]. Brain functional states likely also differ between women and men in the insula; for example, both basal state and stress responses mediated by the HPA-axis show extensive sex-related variations [68].

Existing evidence supports an anterior dominance of the insula during sympathetic regulation [18, 35, 44, 45, 69]. Consistent with this literature, we found that all gyri had a similar response pattern, but relative to the posterior gyrus of each hemisphere (PLG), anterior gyri showed significantly stronger responses during the cold pressor. However, the anterior dominance may also reflect processing of sensory signals, as it has been suggested that the middle insula receives and integrates input relating to bodily feelings from all sensory systems and transfers the signals to the anterior insula, which processes subjective and emotional feelings of the body [70].

Right-sided sympathetic dominance of the insula is a common theme in animal and human studies [12-14, 71-74]. Indeed, in our study, the ASG and MSG showed greater right than left activity throughout the cold pressor phase in both men and women. However, the present study was not well-suited to interpret lateralization effects since it was a right-foot cold pressor, although sex differences in lateralization of the current results can still be interpreted. Sex differences in laterality emerged transiently in the PSG, ALG and PLG, suggest some variation between the sexes in the left-right functional organization of the insula. It is possible that the findings indeed reflect lateralized function but further studies are needed to verify this hypothesis. Using the handgrip task, the right insula shows greater activity than the left, irrespective of which hand is being used during sympathetic phases [75]. It has recently been suggested that the asymmetric processing patterns of autonomic regions like the insula are based on lateralization (ipsi- vs. contralateral) of descending neural pathways that control specific organs [73]. Overall, sympathetic control of the heart is thought to be right-lateralized (including a variety of non-insular brain regions), but the heart receives both sympathetic and parasympathetic input and the slowing of the heart rate can result from either an increase in parasympathetic, or a decrease in sympathetic influences [76]. Visual stimulation of the left hemisphere evoked higher parasympathetic output (vagal tone assessed using the high-frequency component of the ECG-measured heart rate variability) than right visual stimulation. In a separate study, visual stimulation of the right hemisphere increased cardiac output without causing changes in heart rate [77]. The mechanism these latter studies targeted did not engage the cardiovascular system directly, but instead engaged sensory pathways via the limbic system. However, it appears that autonomic output can be generated via different pathways, whereby lateralization patterns remain consistently right-sympathetic and left-parasympathetic. In other words, left and right insular cortices appear to have specialized tasks independent of which side of the body is stimulated. It is therefore possible that the findings do reflect right-sympathetic dominance, not just right-foot stimulation.

Limitations of the current study include a lack of assessment of menstrual cycle and menopausal status, which can influence response elicited by the cold pressor, as well as other autonomic functions [78–85]. Similarly, it is possible that the experience of cold, including cold pain, is systematically different between men and women, and that the fMRI differences may reflect pain rather than cardiovascular differences. Future studies could benefit from asking participants to rate the pain level of the cold pressor. Another limitation is that our sample included several left-handed participants. Handedness has been associated with variations in autonomic markers including the QRS-T interval [86] and muscle sympathetic nerve activity (MSNA) responses during the hand grip [87]. The segmentation by gyri may miss important functional anatomy, including inferior-superior distinctions given the cytoarchitectonic gradient in the insula from inferior (agranular cortex) in the anterior region over the middle (dysgranular cortex) to the posterior (granular cortex) [88–90]. Despite these limitations, our results inspire further research into the details of the role of the insula in sex differences of autonomic challenge responses.

In conclusion, the insula shows sex differences in its functional topography during the sympathetic cold pressor challenge, and the differences are primarily expressed in the right anterior insula, which is consistent with other pressor autonomic challenges we previously investigated. The findings reinforce the role that the right anterior insula plays in autonomic regulation, as well as task-related sensory perception. It is possible that the sex differences in insular responses to the cold pressor and other challenges may be associated with sex differences in physiology underlying cardiovascular risk and disease. These outcomes might be directly or indirectly associated with sensitivity to thermal stress and injuries, autonomic responsiveness and resilience. Sex differences in cardiovascular functioning, stress, pain tolerance, cold resistance and body composition may interact in a complex pattern to lead to the sex differences in insular gyri responses. With regards cardiovascular regulation, although transient male-female differences in fMRI responses appeared during the cold pressor, unlike the Valsalva and handgrip, no sex differences in ASG lateralization emerged, suggesting that the previously found sex differences may not solely relate to regulating blood pressure responses.

## Acknowledgements

None.

## References

1. Ndayisaba JP, Fanciulli A, Granata R, Duerr S, Hintringer F, Goebel G, et al. Sex and age effects on cardiovascular autonomic function in healthy adults. Clin Auton Res. 2015;25(5):317–26. doi: 10.1007/s10286-015-0310-1. PubMed PMID: 26285905.

2. Merz AA, Cheng S. Sex differences in cardiovascular ageing. Heart. 2016;102(11):825–31. doi: 10.1136/heartjnl-2015-308769. PubMed PMID: 26917537.

3. Matsukawa T, Sugiyama Y, Watanabe T, Kobayashi F, Mano T. Gender difference in age-related changes in muscle sympathetic nerve activity in healthy subjects. Am J Physiol. 1998;275(5 Pt 2):R1600–4. PubMed PMID: 9791079.

4. Ng AV, Callister R, Johnson DG, Seals DR. Age and gender influence muscle sympathetic nerve activity at rest in healthy humans. Hypertension. 1993;21(4):498–503. PubMed PMID: 8458648.

5. Koenig J, Thayer JF. Sex differences in healthy human heart rate variability: A meta-analysis. Neurosci Biobehav Rev. 2016;64:288–310. doi: 10.1016/j.neubiorev.2016.03.007. PubMed PMID: 26964804.

6. Liao D, Barnes RW, Chambless LE, Simpson RJ, Jr., Sorlie P, Heiss G. Age, race, and sex differences in autonomic cardiac function measured by spectral analysis of heart rate variability--the ARIC study. Atherosclerosis Risk in Communities. Am J Cardiol. 1995;76(12):906–12. PubMed PMID: 7484830.

7. Abdel-Rahman AR, Merrill RH, Wooles WR. Gender-related differences in the baroreceptor reflex control of heart rate in normotensive humans. J Appl Physiol (1985). 1994;77(2):606–13. PubMed PMID: 8002506.

8. Dart AM, Du XJ, Kingwell BA. Gender, sex hormones and autonomic nervous control of the cardiovascular system. Cardiovasc Res. 2002;53(3):678–87. PubMed PMID: 11861039.

9. Benarroch EE. The central autonomic network: functional organization, dysfunction, and perspective. Mayo Clin Proc. 1993;68(10):988–1001. PubMed PMID: 8412366.

10. Barron SA, Rogovski Z, Hemli J. Autonomic consequences of cerebral hemisphere infarction. Stroke. 1994;25(1):113–6. PubMed PMID: 8266357.

11. Laowattana S, Zeger SL, Lima JA, Goodman SN, Wittstein IS, Oppenheimer SM. Left insular stroke is associated with adverse cardiac outcome. Neurology. 2006;66(4):477–83; discussion 63. doi: 10.1212/01.wnl.0000202684.29640.60. PubMed PMID: 16505298.

12. Oppenheimer SM, Hachinski VC. The cardiac consequences of stroke. Neurologic clinics. 1992;10(1):167–76. Epub 1992/02/01. PubMed PMID: 1557001.

13. Oppenheimer SM, Kedem G, Martin WM. Left-insular cortex lesions perturb cardiac autonomic tone in humans. Clin Auton Res. 1996;6(3):131–40. PubMed PMID: 8832121.

14. Meyer S, Strittmatter M, Fischer C, Georg T, Schmitz B. Lateralization in autonomic dysfunction in ischemic stroke involving the insular cortex. Neuroreport. 2004;15(2):357–61. PubMed PMID: 15076768.

15. Guenot M, Isnard J, Sindou M. Surgical anatomy of the insula. Adv Tech Stand Neurosurg. 2004;29:265–88. doi: 10.1007/978-3-7091-0558-0_7. PubMed PMID: 15035341.

16. Landau E. The Comparative Anatomy of the Nucleus Amygdalae, the Claustrum and the Insular Cortex. J Anat. 1919;53(Pt 4):351–60. PubMed PMID: 17103874; PubMed Central PMCID: PMCPMC1262873.

17. Ture U, Yasargil DC, Al-Mefty O, Yasargil MG. Topographic anatomy of the insular region. J Neurosurg. 1999;90(4):720–33. Epub 1999/04/08. PubMed PMID: 10193618.

18. Oppenheimer S, Cechetto D. The Insular Cortex and the Regulation of Cardiac Function. Compr Physiol. 2016;6(2):1081–133. doi: 10.1002/cphy.c140076. PubMed PMID: 27065176.

19. Shipley MT. Insular cortex projection to the nucleus of the solitary tract and brainstem visceromotor regions in the mouse. Brain Res Bull. 1982;8(2):139–48. PubMed PMID: 7066705.

20. Allen GV, Saper CB, Hurley KM, Cechetto DF. Organization of visceral and limbic connections in the insular cortex of the rat. The Journal of comparative neurology. 1991;311(1):1–16. Epub 1991/09/01. doi: 10.1002/cne.903110102. PubMed PMID: 1719041.

21. Loewy AD. Descending pathways to the sympathetic preganglionic neurons. Prog Brain Res. 1982;57:267–77. Epub 1982/01/01. doi: 10.1016/s0079-6123(08)64133-3. PubMed PMID: 6296919.

22. Nagai M, Hoshide S, Kario K. The insular cortex and cardiovascular system: a new insight into the brain-heart axis. Journal of the American Society of Hypertension : JASH. 2010;4(4):174–82. Epub 2010/07/27. doi: 10.1016/j.jash.2010.05.001. PubMed PMID: 20655502.

23. Cerliani L, Thomas RM, Jbabdi S, Siero JC, Nanetti L, Crippa A, et al. Probabilistic tractography recovers a rostrocaudal trajectory of connectivity variability in the human insular cortex. Hum Brain Mapp. 2012;33(9):2005–34. Epub 2011/07/16. doi: 10.1002/hbm.21338. PubMed PMID: 21761507; PubMed Central PMCID: PMCPMC3443376.

24. Macey PM, Wu P, Kumar R, Ogren JA, Richardson HL, Woo MA, et al. Differential responses of the insular cortex gyri to autonomic challenges. Auton Neurosci. 2012;168(1-2):72–81. Epub 2012/02/22. doi: 10.1016/j.autneu.2012.01.009. PubMed PMID: 22342370; PubMed Central PMCID: PMCPMC4077282.

25. Segerdahl AR, Mezue M, Okell TW, Farrar JT, Tracey I. The dorsal posterior insula subserves a fundamental role in human pain. Nature Neuroscience. 2015;18(4):499–500. doi: 10.1038/nn.3969.

26. Velasco M, Gomez J, Blanco M, Rodriguez I. The cold pressor test: pharmacological and therapeutic aspects. Am J Ther. 1997;4(1):34–8. Epub 1997/01/01. doi: 10.1097/00045391-199701000-00008. PubMed PMID: 10423589.

27. Keller-Ross ML, Cunningham HA, Carter JR. Impact of age and sex on neural cardiovascular responsiveness to cold pressor test in humans. Am J Physiol Regul Integr Comp Physiol. 2020. Epub 2020/07/23. doi: 10.1152/ajpregu.00045.2020. PubMed PMID: 32697654.

28. Miller AJ, Cui J, Luck JC, Sinoway LI, Muller MD. Age and sex differences in sympathetic and hemodynamic responses to hypoxia and cold pressor test. Physiol Rep. 2019;7(2):e13988. Epub 2019/01/20. doi: 10.14814/phy2.13988. PubMed PMID: 30659773; PubMed Central PMCID: PMCPMC6339536.

29. Dishman RK, Nakamura Y, Jackson EM, Ray CA. Blood pressure and muscle sympathetic nerve activity during cold pressor stress: fitness and gender. Psychophysiology. 2003;40(3):370–80. Epub 2003/08/30. doi: 10.1111/1469-8986.00040. PubMed PMID: 12946111.

30. Yamamoto K, Iwase S, Mano T. Responses of muscle sympathetic nerve activity and cardiac output to the cold pressor test. Jpn J Physiol. 1992;42(2):239–52. Epub 1992/01/01. doi: 10.2170/jjphysiol.42.239. PubMed PMID: 1434092.

31. Wirch JL, Wolfe LA, Weissgerber TL, Davies GA. Cold pressor test protocol to evaluate cardiac autonomic function. Appl Physiol Nutr Metab. 2006;31(3):235–43. Epub 2006/06/14. doi: 10.1139/h05-018. PubMed PMID: 16770350.

32. Zhao Q, Bazzano LA, Cao J, Li J, Chen J, Huang J, et al. Reproducibility of blood pressure response to the cold pressor test: the GenSalt Study. Am J Epidemiol. 2012;176 Suppl 7(Suppl 7):S91–8. Epub 2012/10/17. doi: 10.1093/aje/kws294. PubMed PMID: 23035148; PubMed Central PMCID: PMCPMC3530368.

33. Kasagi F, Akahoshi M, Shimaoka K. Relation between cold pressor test and development of hypertension based on 28-year follow-up. Hypertension. 1995;25(1):71–6. Epub 1995/01/01. doi: 10.1161/01.hyp.25.1.71. PubMed PMID: 7843757.

34. Rubenfire M, Rajagopalan S, Mosca L. Carotid artery vasoreactivity in response to sympathetic stress correlates with coronary disease risk and is independent of wall thickness. J Am Coll Cardiol. 2000;36(7):2192–7. Epub 2000/12/29. doi: 10.1016/s0735-1097(00)01021-4. PubMed PMID: 11127460.

35. Richardson HL, Macey PM, Kumar R, Valladares EM, Woo MA, Harper RM. Neural and physiological responses to a cold pressor challenge in healthy adolescents. J Neurosci Res. 2013;91(12):1618–27. Epub 2013/10/10. doi: 10.1002/jnr.23283. PubMed PMID: 24105663; PubMed Central PMCID: PMCPMC4040139.

36. Hutton C, Bork A, Josephs O, Deichmann R, Ashburner J, Turner R. Image distortion correction in fMRI: A quantitative evaluation. Neuroimage. 2002;16(1):217–40. Epub 2002/04/24. doi: 10.1006/nimg.2001.1054 S1053811901910547 [pii]. PubMed PMID: 11969330.

37. Rorden C, Karnath HO, Bonilha L. Improving lesion-symptom mapping. J Cogn Neurosci. 2007;19(7):1081–8. Epub 2007/06/23. doi: 10.1162/jocn.2007.19.7.1081. PubMed PMID: 17583985.

38. Naidich TP, Kang E, Fatterpekar GM, Delman BN, Gultekin SH, Wolfe D, et al. The insula: anatomic study and MR imaging display at 1.5 T. AJNR Am J Neuroradiol. 2004;25(2):222–32. Epub 2004/02/19. PubMed PMID: 14970021.

39. Mavridis I, Boviatsis E, Anagnostopoulou S. Exploring the neurosurgical anatomy of the human insula: a combined and comparative anatomic-radiologic study. Surg Radiol Anat. 2011;33(4):319–28. doi: 10.1007/s00276-010-0699-0. PubMed PMID: 20623284.

40. Littell RC, Milliken GA, Stroup WW, Wolfinger RD. SAS System for Mixed Models. Cary, NC: SAS Institute Inc.; 1996.

41. Macey P, Schluter P, Macey K, Harper R. Detecting variable responses within fMRI time-series of volumes-of-interest using repeated measures ANOVA [version 1; referees: awaiting peer review]. F1000research. 2016;5(563). doi: 10.12688/f1000research.8252.1. PubMed PMID: 10.12688/f1000research.8252.1.

42. Macey PM, Kumar R, Woo MA, Yan-Go FL, Harper RM. Heart rate responses to autonomic challenges in obstructive sleep apnea. PLoS One. 2013;8(10):e76631. Epub 2013/11/07. doi: 10.1371/journal.pone.0076631. PubMed PMID: 24194842; PubMed Central PMCID: PMCPMC3806804.

43. King AB, Menon RS, Hachinski V, Cechetto DF. Human forebrain activation by visceral stimuli. J Comp Neurol. 1999;413(4):572–82. Epub 1999/09/25. doi: 10.1002/(SICI)1096-9861(19991101)413:4<572::AID-CNE6>3.0.CO;2-S [pii]. PubMed PMID: 10495443.

44. Macey PM, Rieken NS, Kumar R, Ogren JA, Middlekauff HR, Wu P, et al. Sex Differences in Insular Cortex Gyri Responses to the Valsalva Maneuver. Front Neurol. 2016;7:87. Epub 2016/07/05. doi: 10.3389/fneur.2016.00087. PubMed PMID: 27375549; PubMed Central PMCID: PMCPMC4899449.

45. Macey PM, Rieken NS, Ogren JA, Macey KE, Kumar R, Harper RM. Sex differences in insular cortex gyri responses to a brief static handgrip challenge. Biol Sex Differ. 2017;8:13. Epub 2017/04/25. doi: 10.1186/s13293-017-0135-9. PubMed PMID: 28435658; PubMed Central PMCID: PMCPMC5397762.

46. Craig AD, Chen K, Bandy D, Reiman EM. Thermosensory activation of insular cortex. Nature neuroscience. 2000;3(2):184–90. Epub 2000/01/29. doi: 10.1038/72131. PubMed PMID: 10649575.

47. Keller-Ross ML, Cunningham HA, Carter JR. Impact of age and sex on neural cardiovascular responsiveness to cold pressor test in humans. Am J Physiol Regul Integr Comp Physiol. 2020;319(3):R288–r95. Epub 2020/07/23. doi: 10.1152/ajpregu.00045.2020. PubMed PMID: 32697654; PubMed Central PMCID: PMCPMC7509253.

48. Zhang M, Zhao Q, Mills KT, Chen J, Li J, Cao J, et al. Factors associated with blood pressure response to the cold pressor test: the GenSalt Study. Am J Hypertens. 2013;26(9):1132–9. Epub 2013/06/04. doi: 10.1093/ajh/hpt075. PubMed PMID: 23727840; PubMed Central PMCID: PMCPMC3741226.

49. Wu T, Snieder H, de Geus E. Genetic influences on cardiovascular stress reactivity. Neurosci Biobehav Rev. 2010;35(1):58–68. Epub 2009/12/08. doi: 10.1016/j.neubiorev.2009.12.001. PubMed PMID: 19963006.

50. Wong SW, Kimmerly DS, Masse N, Menon RS, Cechetto DF, Shoemaker JK. Sex differences in forebrain and cardiovagal responses at the onset of isometric handgrip exercise: a retrospective fMRI study. J Appl Physiol (1985). 2007;103(4):1402–11. Epub 2007/07/07. doi: 10.1152/japplphysiol.00171.2007. PubMed PMID: 17615282.

51. Kimmerly DS, Wong S, Menon R, Shoemaker JK. Forebrain neural patterns associated with sex differences in autonomic and cardiovascular function during baroreceptor unloading. Am J Physiol Regul Integr Comp Physiol. 2007;292(2):R715–22. doi: 10.1152/ajpregu.00366.2006. PubMed PMID: 17272671.

52. Goswami N, Blaber AP, Hinghofer-Szalkay H, Convertino VA. Lower Body Negative Pressure: Physiological Effects, Applications, and Implementation. Physiol Rev. 2019;99(1):807–51. Epub 2018/12/13. doi: 10.1152/physrev.00006.2018. PubMed PMID: 30540225.

53. Uddin LQ. Salience processing and insular cortical function and dysfunction. Nat Rev Neurosci. 2015;16(1):55–61. Epub 2014/11/20. doi: 10.1038/nrn3857. PubMed PMID: 25406711.

54. Craig AD. Topographically organized projection to posterior insular cortex from the posterior portion of the ventral medial nucleus in the long-tailed macaque monkey. J Comp Neurol. 2014;522(1):36–63. doi: 10.1002/cne.23425. PubMed PMID: 23853108; PubMed Central PMCID: PMCPMC4145874.

55. Farrell MJ. Regional brain responses in humans during body heating and cooling. Temperature. 2016;3(2):220–31. doi: 10.1080/23328940.2016.1174794.

56. Davis KD, Kwan CL, Crawley AP, Mikulis DJ. Functional MRI study of thalamic and cortical activations evoked by cutaneous heat, cold, and tactile stimuli. J Neurophysiol. 1998;80(3):1533–46. Epub 1998/09/24. doi: 10.1152/jn.1998.80.3.1533. PubMed PMID: 9744957.

57. Kurth F, Zilles K, Fox PT, Laird AR, Eickhoff SB. A link between the systems: functional differentiation and integration within the human insula revealed by meta-analysis. Brain Struct Funct. 2010;214(5-6):519–34. Epub 2010/06/01. doi: 10.1007/s00429-010-0255-z. PubMed PMID: 20512376; PubMed Central PMCID: PMCPMC4801482.

58. Ruigrok AN, Salimi-Khorshidi G, Lai MC, Baron-Cohen S, Lombardo MV, Tait RJ, et al. A meta-analysis of sex differences in human brain structure. Neurosci Biobehav Rev. 2014;39:34–50. doi: 10.1016/j.neubiorev.2013.12.004. PubMed PMID: 24374381; PubMed Central PMCID: PMCPMC3969295.

59. Rosen A, Chen DQ, Hayes DJ, Davis KD, Hodaie M. A Neuroimaging Strategy for the Three-Dimensional in vivo Anatomical Visualization and Characterization of Insular Gyri. Stereotact Funct Neurosurg. 2015;93(4):255–64. doi: 10.1159/000380826. PubMed PMID: 26066396.

60. Herting MM, Maxwell EC, Irvine C, Nagel BJ. The impact of sex, puberty, and hormones on white matter microstructure in adolescents. Cereb Cortex. 2012;22(9):1979–92. doi: 10.1093/cercor/bhr246. PubMed PMID: 22002939; PubMed Central PMCID: PMCPMC3412439.

61. Hu Y, Xu Q, Shen J, Li K, Zhu H, Zhang Z, et al. Small-worldness and gender differences of large scale brain metabolic covariance networks in young adults: a FDG PET study of 400 subjects. Acta Radiol. 2015;56(2):204–13. doi: 10.1177/0284185114529106. PubMed PMID: 24763919.

62. Saleh TM, Connell BJ. Role of oestrogen in the central regulation of autonomic function. Clin Exp Pharmacol Physiol. 2007;34(9):827–32. doi: 10.1111/j.1440-1681.2007.04663.x. PubMed PMID: 17645624.

63. Saleh TM, Connell BJ, Legge C, Cribb AE. Estrogen attenuates neuronal excitability in the insular cortex following middle cerebral artery occlusion. Brain Res. 2004;1018(1):119–29. doi: 10.1016/j.brainres.2004.05.074. PubMed PMID: 15262213.

64. Hwang MJ, Zsido RG, Song H, Pace-Schott EF, Miller KK, Lebron-Milad K, et al. Contribution of estradiol levels and hormonal contraceptives to sex differences within the fear network during fear conditioning and extinction. BMC Psychiatry. 2015;15(1):295. doi: 10.1186/s12888-015-0673-9. PubMed PMID: 26581193; PubMed Central PMCID: PMCPMC4652367.

65. Saleh MC, Connell BJ, Saleh TM. Resveratrol induced neuroprotection is mediated via both estrogen receptor subtypes, ER(alpha) and ER(beta). Neurosci Lett. 2013;548:217–21. doi: 10.1016/j.neulet.2013.05.057. PubMed PMID: 23748073.

66. Gur RC, Gur RE, Obrist WD, Hungerbuhler JP, Younkin D, Rosen AD, et al. Sex and handedness differences in cerebral blood flow during rest and cognitive activity. Science. 1982;217(4560):659–61. PubMed PMID: 7089587.

67. Rodriguez G, Warkentin S, Risberg J, Rosadini G. Sex differences in regional cerebral blood flow. J Cereb Blood Flow Metab. 1988;8(6):783–9. doi: 10.1038/jcbfm.1988.133. PubMed PMID: 3192645.

68. Goel N, Workman JL, Lee TT, Innala L, Viau V. Sex differences in the HPA axis. Compr Physiol. 2014;4(3):1121–55. doi: 10.1002/cphy.c130054. PubMed PMID: 24944032.

69. Beissner F, Meissner K, Bar KJ, Napadow V. The autonomic brain: an activation likelihood estimation meta-analysis for central processing of autonomic function. J Neurosci. 2013;33(25):10503–11. doi: 10.1523/JNEUROSCI.1103-13.2013. PubMed PMID: 23785162; PubMed Central PMCID: PMCPMC3685840.

70. Craig AD. Significance of the insula for the evolution of human awareness of feelings from the body. Ann N Y Acad Sci. 2011;1225:72–82. Epub 2011/05/04. doi: 10.1111/j.1749-6632.2011.05990.x. PubMed PMID: 21534994.

71. Sykora M, Diedler J, Turcani P, Hacke W, Steiner T. Baroreflex: a new therapeutic target in human stroke? Stroke. 2009;40(12):e678–82. doi: 10.1161/STROKEAHA.109.565838. PubMed PMID: 19834010.

72. Hilz MJ, Dutsch M, Perrine K, Nelson PK, Rauhut U, Devinsky O. Hemispheric influence on autonomic modulation and baroreflex sensitivity. Ann Neurol. 2001;49(5):575–84. PubMed PMID: 11357947.

73. Xavier CH, Mendonça MM, Marins FR, da Silva ES, Ianzer D, Colugnati DB, et al. Stating asymmetry in neural pathways: methodological trends in autonomic neuroscience. Int J Neurosci. 2018;128(11):1078–85. Epub 2018/05/05. doi: 10.1080/00207454.2018.1473396. PubMed PMID: 29724119.

74. Oppenheimer SM, Gelb A, Girvin JP, Hachinski VC. Cardiovascular effects of human insular cortex stimulation. Neurology. 1992;42(9):1727–32. Epub 1992/09/01. doi: 10.1212/wnl.42.9.1727. PubMed PMID: 1513461.

75. Wong SW, Masse N, Kimmerly DS, Menon RS, Shoemaker JK. Ventral medial prefrontal cortex and cardiovagal control in conscious humans. Neuroimage. 2007;35(2):698–708. Epub 2007/02/13. doi: S1053-8119(06)01206-7 [pii] 10.1016/j.neuroimage.2006.12.027. PubMed PMID: 17291781.

76. Wittling W, Block A, Genzel S, Schweiger E. Hemisphere asymmetry in parasympathetic control of the heart. Neuropsychologia. 1998;36(5):461–8. Epub 1998/08/12. doi: 10.1016/s0028-3932(97)00129-2. PubMed PMID: 9699952.

77. Wittling W, Block A, Schweiger E, Genzel S. Hemisphere Asymmetry in Sympathetic Control of the Human Myocardium. Brain and Cognition. 1998;38(1):17–35. doi: https://doi.org/10.1006/brcg.1998.1000.

78. Mogil JS. Sex differences in pain and pain inhibition: multiple explanations of a controversial phenomenon. Nat Rev Neurosci. 2012;13(12):859–66. doi: 10.1038/nrn3360. PubMed PMID: 23165262.

79. Mogil JS. Qualitative sex differences in pain processing: emerging evidence of a biased literature. Nature Reviews Neuroscience. 2020;21(7):353–65. doi: 10.1038/s41583-020-0310-6.

80. Kowalczyk WJ, Evans SM, Bisaga AM, Sullivan MA, Comer SD. Sex differences and hormonal influences on response to cold pressor pain in humans. J Pain. 2006;7(3):151–60. Epub 2006/03/07. doi: 10.1016/j.jpain.2005.10.004. PubMed PMID: 16516820.

81. Stening K, Eriksson O, Wahren L, Berg G, Hammar M, Blomqvist A. Pain sensations to the cold pressor test in normally menstruating women: comparison with men and relation to menstrual phase and serum sex steroid levels. Am J Physiol Regul Integr Comp Physiol. 2007;293(4):R1711–6. Epub 2007/07/27. doi: 10.1152/ajpregu.00127.2007. PubMed PMID: 17652363.

82. Riley JL, 3rd, Robinson ME, Wise EA, Price D. A meta-analytic review of pain perception across the menstrual cycle. Pain. 1999;81(3):225–35. Epub 1999/08/04. doi: 10.1016/s0304-3959(98)00258-9. PubMed PMID: 10431710.

83. Sullivan JM, Shala BA, Miller LA, Lerner JL, McBrayer JD. Progestin enhances vasoconstrictor responses in postmenopausal women receiving estrogen replacement therapy. Menopause. 2018;25(11):1180–6. Epub 2018/10/26. doi: 10.1097/gme.0000000000001214. PubMed PMID: 30358710.

84. Vallejo M, Marquez MF, Borja-Aburto VH, Cardenas M, Hermosillo AG. Age, body mass index, and menstrual cycle influence young women’s heart rate variability --a multivariable analysis. Clin Auton Res. 2005;15(4):292–8. doi: 10.1007/s10286-005-0272-9. PubMed PMID: 16032384.

85. Lavi S, Nevo O, Thaler I, Rosenfeld R, Dayan L, Hirshoren N, et al. Effect of aging on the cardiovascular regulatory systems in healthy women. Am J Physiol Regul Integr Comp Physiol. 2007;292(2):R788–93. doi: 10.1152/ajpregu.00352.2006. PubMed PMID: 16946083.

86. Iscen S, Ozenc S, Tavlasoglu U. Association between left-handedness and cardiac autonomic function in healthy young men. Pacing Clin Electrophysiol. 2014;37(7):884–8. doi: 10.1111/pace.12365. PubMed PMID: 24697725.

87. Saito M, Kato M, Mano T. Comparison of sympathetic nerve activity during handgrip exercise performed with the dominant and non-dominant arm. Environ Med. 2000;44(1):60–2. PubMed PMID: 12296372.

88. Mesulam MM, Mufson EJ. Insula of the old world monkey. I. Architectonics in the insulo-orbito-temporal component of the paralimbic brain. J Comp Neurol. 1982;212(1):1–22. doi: 10.1002/cne.902120102. PubMed PMID: 7174905.

89. Morel A, Gallay MN, Baechler A, Wyss M, Gallay DS. The human insula: Architectonic organization and postmortem MRI registration. Neuroscience. 2013;236:117–35. doi: 10.1016/j.neuroscience.2012.12.076. PubMed PMID: 23340245.

90. Klein TA, Ullsperger M, Danielmeier C. Error awareness and the insula: links to neurological and psychiatric diseases. Front Hum Neurosci. 2013;7:14. doi: 10.3389/fnhum.2013.00014. PubMed PMID: 23382714; PubMed Central PMCID: PMCPMC3563042.

